# Efficiency Estimates for Electromicrobial Production of Branched-chain Hydrocarbons

**DOI:** 10.1101/2023.03.03.531000

**Authors:** Timothy J. Sheppard, David A. Specht, Buz Barstow

**Author notes:** Corresponding author: Buz Barstow, 228 Riley-Robb Hall, Cornell University, Ithaca, NY 14853.

## Abstract

Electromicrobial production is a process where microorganisms use electricity as a charge and energy source for the production of complex molecules, often from starting compounds as simple as CO_2_. The aviation industry is in need for sustainable fuel alternatives that can meet their requirements of high-altitude performance while also meeting 21^st^ century carbon emissions standards. The electromicrobial production of jet fuel components with CO_2_-derived carbon provides a unique opportunity to generate jet fuel blends that are compatible with modern engines with net-neutral carbon emissions. In this study, we analyze the pathways necessary to generate single- and multi-branched-chain hydrocarbons *in vivo* utilizing both extracellular electron uptake (EEU) and H_2_-oxidation as methods for electron delivery, the Calvin cycle for CO_2_-fixation and the ADO decarboxylation pathway. We find the maximum electrical-to-fuel energy conversion efficiencies for single- and multi-branched chain hydrocarbons are 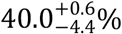 and 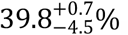. Utilizing this information, as well as previously collected predictions on straight-chain alkane and terpenoid biosynthesis, we calculate the efficiency of electromicrobial production of jet fuel blends containing straight-chain, branched-chain, and terpenoid components. Increasing the fraction of branched-chain alkanes in the blend from zero to 47% only lowers the electrical energy conversion efficiency from 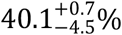 to 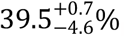.

## Introduction

The industrial-scale synthesis of carbon-neutral hydrocarbon fuels that are drop-in compatible with present-day internal combustion and jet engines is one of the biggest challenges in decarbonization of world’s energy infrastructure. Corn ethanol can be safely blended into gasoline up to a fraction of ≈ 10-15%. However, the widespread use of high fractions of ethanol in fuel blends is unfeasible as it lacks the high boiling and low freezing points needed in many applications, especially aviation^1^. This difficulty has motivated the production of kerosene-grade biofuel blends. Third-generation algal biofuels show promise for use in aviation, but the upscaling of algal fuels is challenging and faces many hurdles to commercial acceptance^2^. For alternative fuels to be established as commonplace, their properties must be as close to those of conventional fuels as possible.

In addition to straight-chain alkanes and terpenoids, branched-chain hydrocarbons are key components of traditional jet fuels. In Jet A1 branched-chain hydrocarbons are used to raise the boiling, and lower the freezing points while burning almost as cleanly as straight-chain alkanes^3^. Although terpenoids (whose electromicrobial synthesis was described by us previously^4^) could achieve similar boiling increases and freezing point reductions, they also create significant soot deposition during combustion^5^. Production of a library of branched-chain hydrocarbons could permit synthesis of blends that closely match the composition and physico-chemical properties of fossil-derived kerosene, while also burning cleanly. Furthermore, gasoline containing a high fraction of isoalkanes (one type of branched-chain alkanes) could burn much cleaner than conventional gasoline^6^.

Electromicrobial production (EMP) could enable highly efficient production of carbon-neutral drop-in biofuels. EMP is a broadly-encompassing term for a group of technologies that aim to combine electricity and microbial metabolism for conversion of simple molecules like CO_2_, CO, HCOO^-^, and N_2_ into complex, energy dense molecules like food and biofuels^7-16^. EMP includes technologies like microbes that assimilate electrochemically-reduced CO_2_ like formate^14,17^; H_2_-oxidizing, CO_2_-fixing systems like the Bionic Leaf^18,19^; microbe-semiconductor hybrids^20^; and microbes that can directly absorb electricity through processes like extracellular electron uptake (EEU)^8,21,22^. Lab-scale demonstrations of EMP already have effective solar-to-chemical energy conversion efficiencies exceeding all forms of terrestrial photosynthesis^19,23^. Meanwhile, theoretical predictions indicate that the efficiency of EMP could exceed all forms of photosynthesis^7,8,24-26^. This high efficiency mitigates many of the concerns about competition for land created by first- and second-generation biofuels^27,28^. Furthermore, a large library of metabolic pathways for the biological synthesis of branched-chain hydrocarbons has been established^3,6,29-32^ that could allow the production of jet fuel blends much closer in composition to Jet A1 than algae-derived biofuels (reviewed in Adesina *et al*.^33^ and Sheppard *et al*.^4^).

In this work we extend our earlier predictions of electromicrobial production efficiency^4,8,25,26^ to make minimum energy cost and upper-limit production efficiency estimates of single- and multi-branched hydrocarbons powered by H_2_-oxidation^18,19^ or EEU^8,21,22^, with carbon supplied by *in vivo* CO_2_-fixation with the Calvin cycle. We then calculate the production efficiency of drop-in fuel blends of increasing branched-chain content.

## Theory

### Electromicrobial Production of Jet Fuel Components

We predict upper limit efficiencies for the electromicrobial production of branched-chain hydrocarbons. These predictions set an upper bound on the performance of a set of highly-engineered microorganisms created for production of drop-in jet fuel components. Below we summarize all of the key equations utilized in this article. For detailed derivations see Salimijazi *et al*.^8^ and our subsequent work that builds upon this theory^4,25,26^. All model parameters are shown in **Table 1**, and all symbols used in this article are shown in **Table S1**.

**Table 1.**
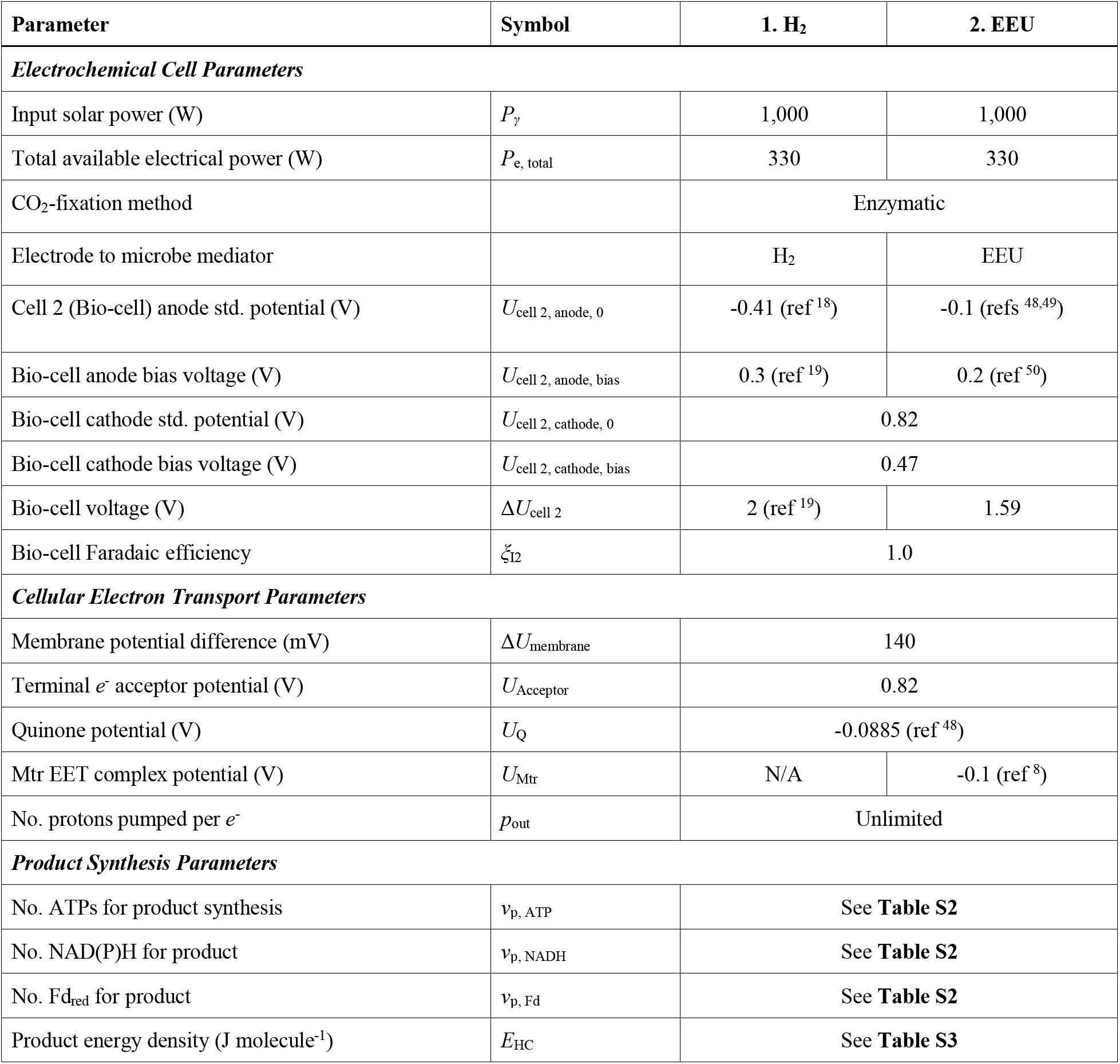
Electromicrobial jet fuel production model parameters. Model parameters used in this article are based upon model parameters used in a previous analysis of the electromicrobial production of the biofuel butanol^8^. A sensitivity analysis was performed for all key parameters in this work^8^.

As in earlier work, we assume access to a reservoir of CO_2_. Reducing power for the regeneration of NAD(P)H and ATP are provided via oxidation of electrochemically-reduced H_2_ (**Fig. 1B** part 1) or through by EEU from a diffusible intermediary (such as flavins or anthra(hydra)quinone-2,6-disulfonate (AHDS_red_/AQDS_ox_)) or through a conductive biofilm or direct contact with a cathode (**Fig. 1B** part 2). We assume that the energy requirements for microbial maintenance are negligible at maximum efficiency, allowing the cell to operate as a “bag of enzymes”(refs ^8,34^).

**Figure 1.**
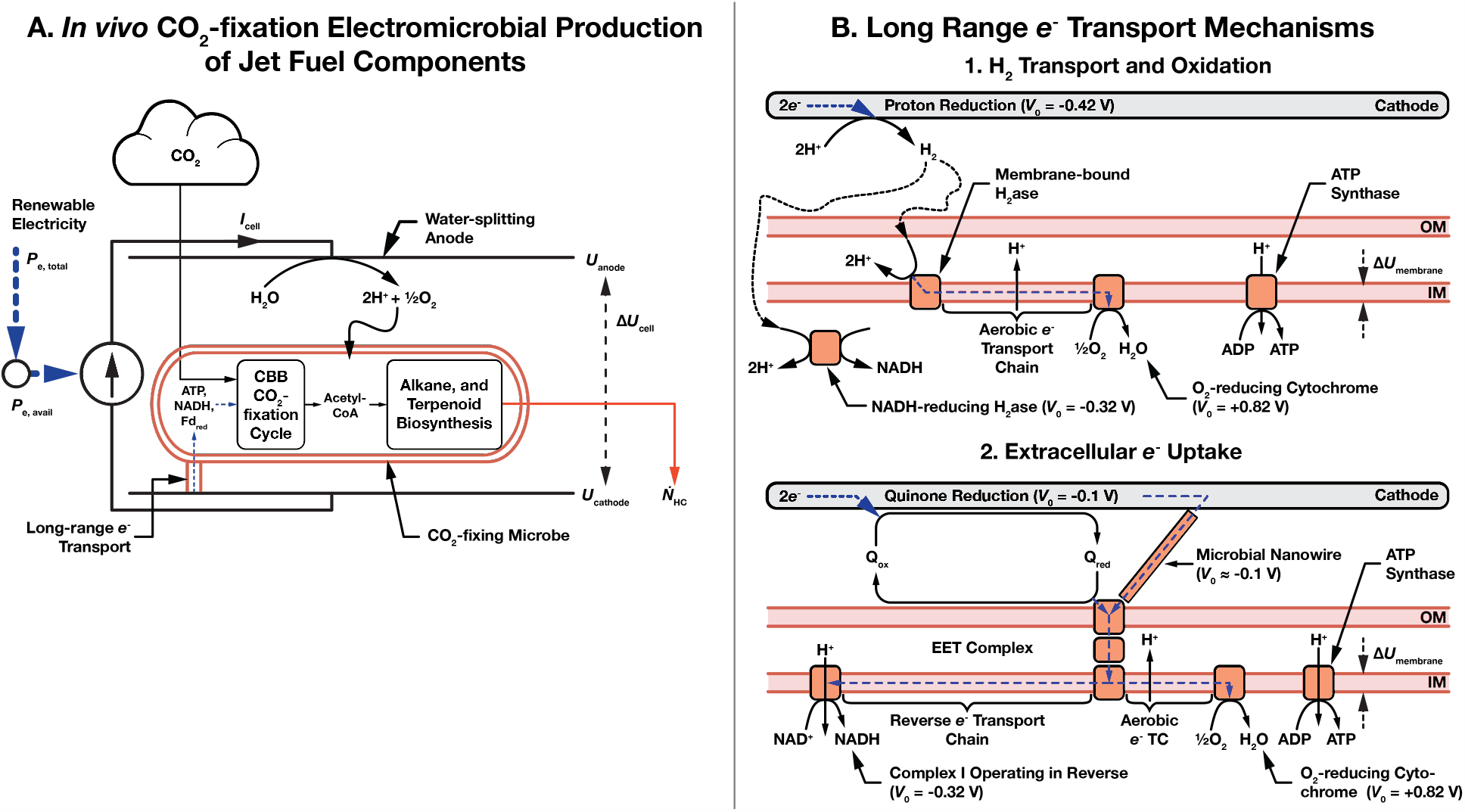
Schematic of electromicrobial production of jet fuel components. **(A)** In the article we just consider electromicrobial production systems that use the Calvin-Benson-Bassham (CBB) cycle for *in vivo* CO_2_-fixation and hydrocarbon synthesis. **(B)** Mechanisms by which electricity sources can be used to power microbial production, using either H_2_-oxidation or extracellular electron uptake (EEU). In the first, H_2_ is electrochemically reduced on a cathode, transferred to the microbe by diffusion or stirring, and is enzymatically oxidized. In the second mechanism, extracellular electron uptake (EEU), electrons are transferred from a cathode (*i*) along a microbial nanowire (part of a conductive biofilm), or (*ii*) by a reduced medium potential redox shuttle like a quinone or flavin, or (*iii*, not shown) by direct contact of the cell with the cathode, and are then oxidized at the cell surface by the extracellular electron transfer (EET) complex. From the thermodynamic perspective considered in this article, these mechanisms of EEU are equivalent. Electrons are then transported to the inner membrane where reverse electron transport is used to regenerate NAD(P)H, reduced Ferredoxin (not shown), and ATP^8,21,42^. This schematic is modified from our earlier work on the synthesis of the straight-chain alkane and terpenoid components of jet fuels^4^.

Hydrocarbon molecules with an energy-per-molecule, *E*_HC_, are produced a rate of 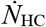 molecules per second. The amount of energy needed to produce a mole of hydrocarbon, *L*_EP_ (refs ^8^ and ^25^),

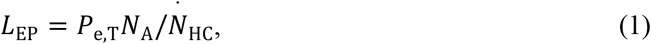

where *N*_A_ is the Avogadro constant. Thus, the minimum energy input into the bio-electrochemical system is,

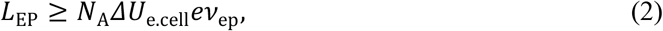

where Δ*U*_e.cell_ is the potential difference across the bio-electrochemical cell (note we have changed this from Δ*U*_cell_ in earlier work for clarity), *e* is the fundamental charge, and *ν*_ep_ is the number of electrons needed to synthesize a molecule of the product from CO_2_.

Furthermore, the efficiency of energy conversion from input power to final product is,

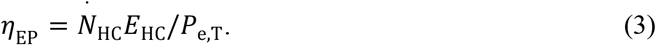

When utilizing *in vivo* carbon-fixation (**Fig. 1A**), the upper limit of electrical-to-chemical efficiency is equivalent to the energy carried per molecule of hydrocarbon, *E*_HC_, relative to the amount of energy needed to move the charge for product synthesis across the bio-electrochemical cell (*eν*_ep_Δ*U*_e.cell_)^8^,

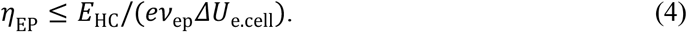

In this article we calculate the number of electrons needed for production of a hydrocarbon by *in vivo* CO_2_-fixation (*ν*_ep_) with electron uptake both by H_2_-oxidation and EEU^8^. For electron delivery by H_2_,

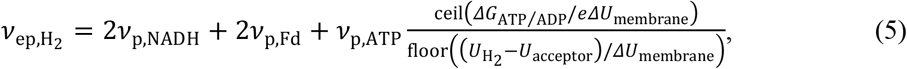

where *ν*_p, NADH_, *ν*_p, Fd_, and *ν*_p, ATP_ are the number of NAD(P)H, reduced ferredoxin, and ATP needed for product synthesis; Δ*G*_ATP/ADP_ is the Gibbs free energy for regeneration of ATP; Δ*U*_membrane_ is the potential difference between the cytoplasmic and periplasmic faces of the inner membrane (the host of the electron transport chain); *U*_H2_ is the redox potential of H_2_-oxidation; and *U*_acceptor_ is the redox potential of the terminal electron acceptor (usually O_2_).

For electron delivery by EEU,

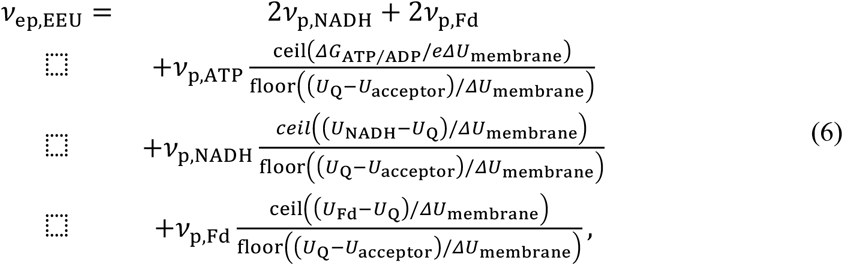

where *U*_NADH_ is the redox potential of NAD(P)H reduction; *U*_Q_ is the redox potential of menaquinone reduction; and *U*_Fd_ is the redox potential of ferredoxin reduction.

### ATP, NAD(P)H, and Reduced Ferredoxin Demands for Jet Fuel Component Electromicrobial Production

The ATP, NADP(H), and reduced ferredoxin requirements for individual molecules in a jet fuel blend are calculated by flux balance analysis. Pathways for the production of the straight-chain alkane and terpenoid components of jet fuel were compiled by us in a recent article^4^.

Like straight-chain alkanes, branched-chain alkanes are produced by the Type II Fatty Acid Synthesis (FAS) system followed by decarboxylation^3^. Branches are introduced into the growing fatty acid by incorporation of unconventional methylated initiator and lengthener molecules^3^. An overview of branched-chain alkane synthesis is shown in **Fig. 2**. Synthesis pathways for methylated initiators and lengtheners are shown in **Fig. 3** and **Table 2**. In wild-type cells, the incorporation of methylated initiators and lengtheners into fatty acids is kept at low levels, and some aspects of these systems are down-regulated by one of several regulatory enzymes native to these systems^3,35^. Down-regulation of these enzymes can promote methylated initiator production and a high output of branched-chain hydrocarbons. From this start point, we are able to generate single-branched compounds of any length with odd or even methylation patterns as shown in **Fig. 3**. Full pathways for the synthesis of a panel of individual branched-chain alkane compounds (shown in **Fig. 4**) are compiled from listings of reactions in the Kyoto Encyclopedia of Genes and Genomes (KEGG)^36-38^ in the input files to the INFO-FIG4A&B.PY, INFO-FIG4C&D.PY codes in the EMP-TO-BRANCHED-JET repository^39^.

**Table 2.** Reactions for synthesis of initiator and lengthener molecules used for branched-chain alkane production. Full pathways collected from the Kyoto Encyclopedia of Genes and Genomes^36-38^. The (S)-methyl-malonyl-[acp] pathway is mediated by the down-regulation of a naturally-occurring regulatory enzyme MCoAD (methylmalonyl-CoA Demethylase) that traditionally prevents (S)-methylmalonyl-CoA formation.

| No. | Reaction | E.C. Number | KEGG Accession Code |
| --- | --- | --- | --- |
|  | **(S)-methyl-malonyl-[acp]** |  |  |
| 1 | ATP + propionyl-CoA + HCO<sub>3</sub><sup>-</sup> → ADP + orthophosphate + (S)-methylmalonyl-CoA | 2.3.1.39 | R00742 |
| 2 | (S)-methylmalonyl-CoA + H-[acp] → (S)-methylmalonyl—[acp] + H-CoA | 2.3.1.39 | R01626 |
|  | **2-methyl-propionyl—[acp]** |  |  |
| 1 | ATP + acetyl-CoA + HCO<sub>3</sub><sup>-</sup> → ADP + orthophosphate + malonyl-CoA | 2.3.1.39 | R00742 |
| 2 | Malonyl-CoA + H-[acp] → malonyl-[acp] + H-CoA | 2.3.1.39 | R01626 |
| 3 | Acetyl-CoA + H-[acp] → acetyl-[acp] + H-CoA | 2.3.1.86 | R01624 |
| 4 | Acetyl-[acp] + malonyl-[acp] → acetoacetyl-[acp] + CO<sub>2</sub> + CoA | 2.3.1.41 | R04355 |
| 5 | Acetoacetyl-[acp] + NADPH + H<sup>+</sup> → 3R-3-hydroxybutanoyl-[acp] + NADP<sup>+</sup> | 1.1.1.100 | R02767 |
| 6 | 3R-3-hydroxybutanoyl-[acp] → but-2-enoyl-[acp] + H<sub>2</sub>O | 4.2.1.59 | R04428 |
| 7 | But-2-enoyl-[acp] + NADPH + H<sup>+</sup> → butanoyl-[acp] + NADP<sup>+</sup> | 1.3.1.9 | R04429 |
| 8 | Butanoyl-[acp] → 2-methyl-propanoyl-[acp] | 5.4.99.3 | R03052 |
|  | **3-methyl-butanoyl—[acp]** |  |  |
| 1 | 2 × pyruvate → (S)-2-acetolactate + CO<sub>2</sub> | 2.2.1.6 | R00006 |
| 2 | (S)-2-acetolactate → 3-hydroxy-3-methyl-2-oxobutanoic acid | 1.1.1.86 | R04439 |
| 3 | (R)-2,3-Dihydroxy-3-methylbutanoate → 3-methyl-2-oxobutanoic acid + H<sub>2</sub>O | 4.2.1.9 | R01209 |
| 4 | Acetyl-CoA + 3-Methyl-2-oxobutanoic acid + H<sub>2</sub>O → α-isopropylmalate + CoA | 2.3.3.13 | R01213 |
| 5 | α-isopropylmalate → 2-isopropylmalate + H<sub>2</sub>O | 4.2.1.33 | R03968 |
| 6 | 2-isopropylmalate + H<sub>2</sub>O → (2R,3S)-3-isopropylmalate | 4.2.1.33 | R04001 |
| 7 | (2R,3S)-3-isopropylmalate + NAD<sup>+</sup> → (2S)-2-isopropyl-3-oxosuccinate + NADH + H<sup>+</sup> | 1.1.1.85 | R01652 |
| 8 | (2S)-2-isopropyl-3-oxosuccinate → 4-methyl-2-oxopentanoate + CO<sub>2</sub> | 1.1.1.85 | R01652 |
| 12 | 4-methyl-2-oxopentanoate + CoA + NAD<sup>+</sup> → 3-methyl-butanoyl-CoA + CO<sub>2</sub> + NADH + H<sup>+</sup> | 1.2.1.25 | R01651 |
| 12 | $3\text{-methyl-butanoyl-CoA} + \text{H-[acp]} \rightarrow 3\text{-methyl-butanoyl-[acp]} + \text{H-CoA}$ | 2.3.1.39 | R01626 |

**Figure 2.**
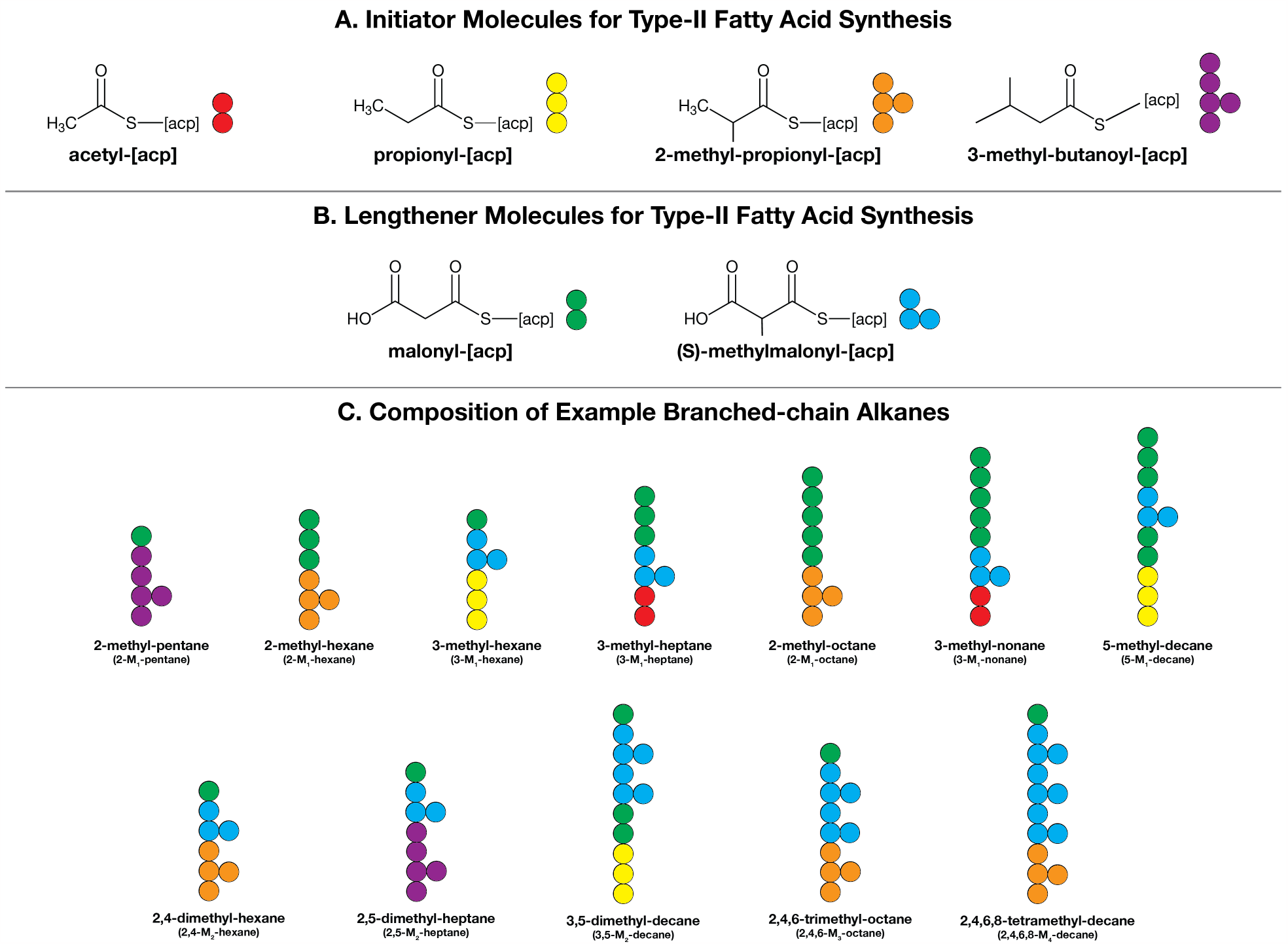
Mechanics for branched-chain alkane production. Branched-chain alkanes are synthesized by the same type-II fatty acid synthesis (FAS) system as straight-chain alkanes^4^, but branches are added with additional initiator and lengthener molecules. (**A**) Initiators for branched-chain alkane synthesis. Acetyl-[acp] (acp: acyl-carrier protein) and propionyl-[acp] are also used as initiators for synthesis of even and odd chain-length straight-chain alkanes by type II fatty acid synthesis^4^. 2-methyl-propionyl-[acp] and 3-methyl-butanoyl-[acp] are used exclusively for branched-chain alkanes by type II fatty acid synthesis. (**B**) Lengtheners for branched-chain alkane synthesis. Malonyl-[acp] is used to add two additional carbons to a growing straight-or branched-chain alkane. (S)-methyl-malonyl-[acp] is used to add a branch to a growing branched-chain alkane. In all cases considered in this article, the last carbon in the alkane is by a termination reaction catalyzed by the well Aldehyde Deformolating Oxygenase (ADO) pathway. (**C**) Composition of example branched-chain alkane molecules shown in **Figure 4**. Synthesis pathways for initiators and lengtheners are shown in Figure 3. Abridged pathways are shown in **Table 3**. Note that the position of the branch is normally measured from the bottom (the start of synthesis) of the molecule, but in the case of 3-M_1_-hexane and 5-M_1_-decane it is measured from the top of the chain.

**Figure 3.**
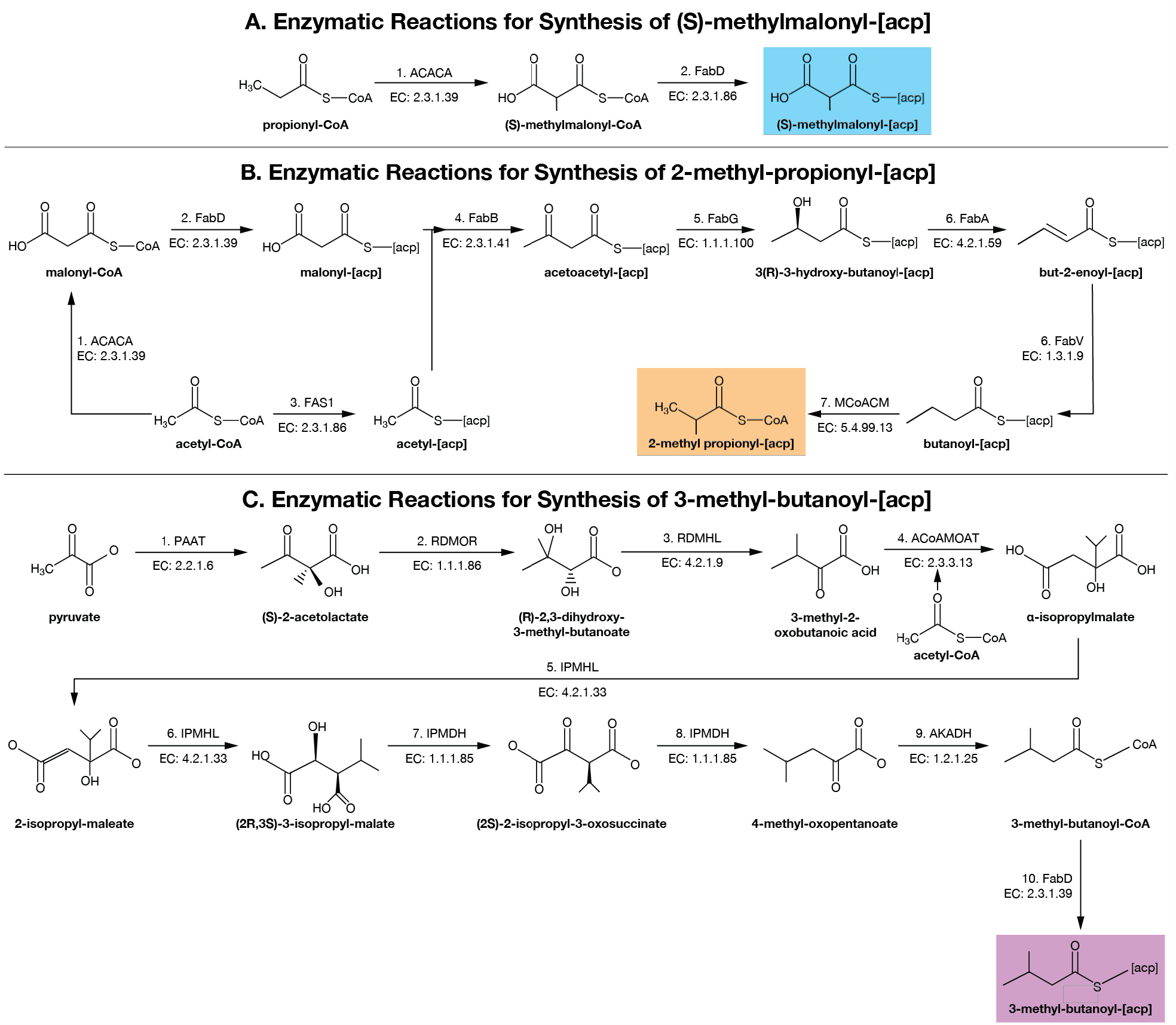
Synthesis pathways for (**A**) the branched chain lenghtener (S)-methylmalonyl-[acp] and branched chain initiators (**B**) 2-methyl-propionyl-[acp] and **(C)** 3-methyl-butanoyl-[acp]. Full pathways collected from Kyoto Encyclopedia of Genes and Genomes^36-38^. (S)-methylmalonyl-[acp] pathway mediated by the down-regulation of a naturally occurring regulatory enzyme MCoAD (methylmalonyl-CoA Demethylase) that traditionally prevents (S)-methylmalonyl-CoA formation. Full pathways list can be found in **Table 2**.

**Figure 4.**
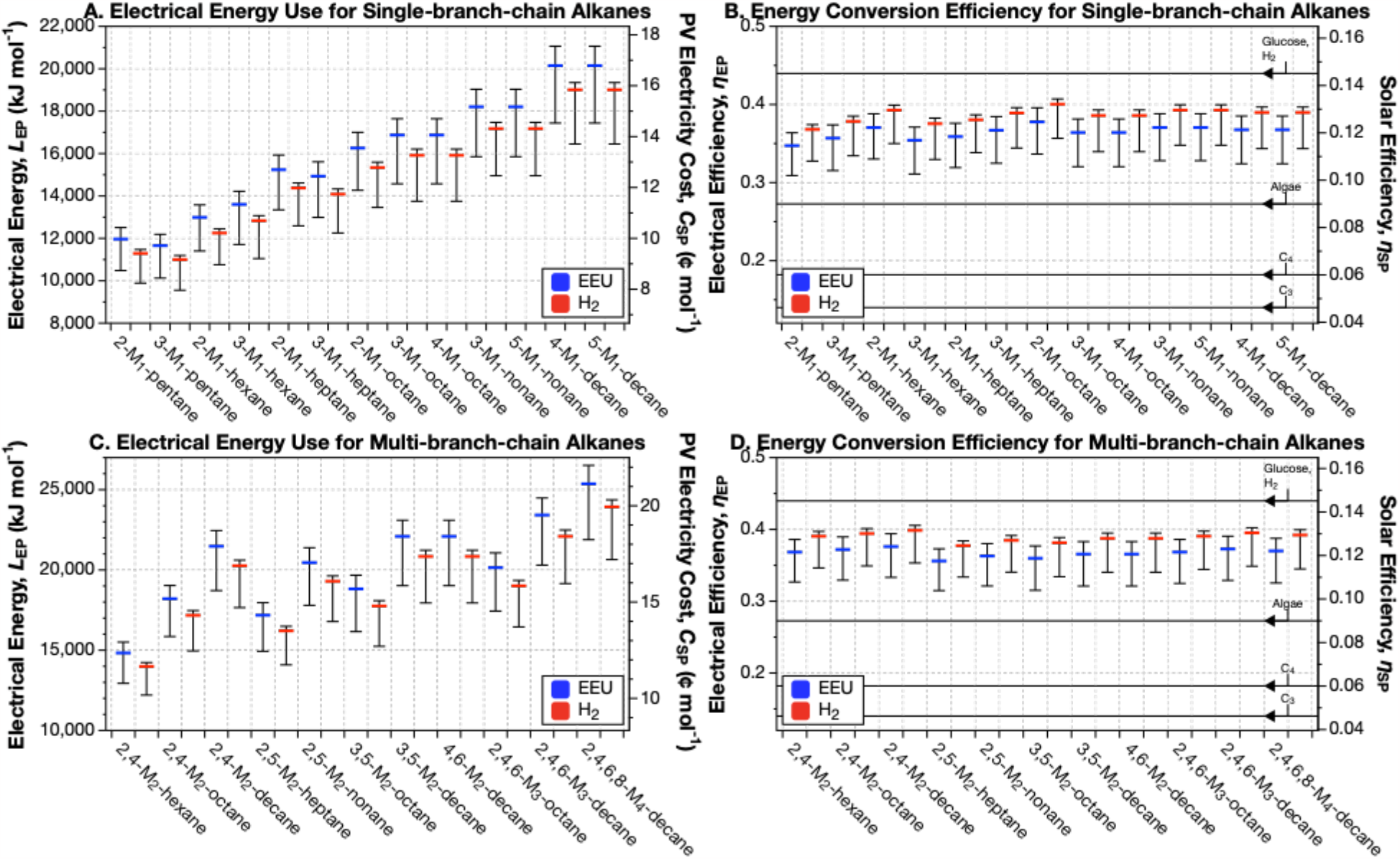
Electrical energy requirements and energy conversion efficiencies for single-and multi-branched-chain alkane production yields maximum efficiencies of 40.0% and 39.9% respectively. (**A**) Energy input for single-branched-chain fatty alkane biosynthesis using the Calvin CO_2_-fixation cycle with the ADO alkane termination pathway. (**B**) Energy conversion efficiency of single-branched-chain from solar cell on left axis. Solar conversion efficiency compared to C_3_, C_4_, Algae, and Glucose on right axis. (**C**) Energy input required for multi-branched-chain alkane compound biosynthesis. (**D**) Energy conversion efficiency of multi-branched-chain alkane compound biosynthesis on left axis. Solar conversion efficiency compared to C_3_, C_4_, Algae, and Glucose on right axis, lines corresponding to those in panel **B**. A sensitivity analysis by Salimijazi *et al*.^8^ found that the biggest source of uncertainty in the energy input and efficiency calculation is the potential difference across the inner membrane of the cell (Δ*U*_membrane_). Estimates for the trans-membrane voltage range from 80 mV (BioNumber ID^43^ (BNID) 10408284 to 270 mV (BNID 107135), with a most likely value of 140 mV (BNIDs 109774, 103386, and 109775). The central value (thick blue or red bar) corresponds to 140 mV. Our sensitivity analysis found that Δ*U*_membrane_ = 280 mV produces lower efficiencies (hence a higher energy input), while Δ*U*_membrane_ = 80 mV produces higher efficiencies (and hence lower energy inputs)^8^. The right axis in panels **A** and **C** shows the minimum cost of that solar electricity, assuming that the United States Department of Energy’s cost target of 3 ¢ per kWh by 2030 can be achieved^44^. The right axes in panels **B** and **D** show the solar-to-product energy conversion efficiency, assuming the system is supplied by a perfectly efficient single-junction Si solar photovoltaic (solar to electrical efficiency of 32.9% (ref ^40^). For comparison, we have marked the upper limit solar-to-biomass energy conversion efficiencies of C_3_, C_4_ (refs ^45,46^), algal photosynthesis^47^, and upper limit electromicrobial production conversion efficiency of glucose using H_2_-oxidation and the Calvin cycle^25^ on the right axes of panels **B** and **D**. This figure can be reproduced by running the codes INFO-FIG-4A&B.PY, INFO-FIG-4C&D.PY, FIG-4A&B.PY, and FIG-4C&D.PY in the EMP-TO-BRANCHED-JET online code repository^39^.

The overall stoichiometric matrix (**S**_**p**_) for synthesis of each alkane using the Calvin-Benson-Bassham cycle was calculated by the INFO-FIG4A&B.PY and INFO-FIG4C&D.PY codes^39^. Briefly, we consider a chemical species number rate of change vector, 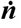, that encodes the rate of change of number of the reactant molecules over a single cycle of the reaction network; a stoichiometric matrix **S**_p_ that encodes the number of reactants made or consumed in every reaction in the network; and a flux vector ***v*** that encodes the number of times each reaction is used in the network. Reactant molecules are denoted as inputs (*e*.*g*., CO_2_, ATP, NAD(P)H), outputs (*e*.*g*., H_2_O), intermediates, or the target molecule (*e*.*g*., the alkane to be synthesized). For the purposes of this thermodynamic analysis, we consider NADH and NADPH to be equivalent as they have near identical redox potentials. The number of NAD(P)H, reduced Ferredoxin and ATP for each individual alkane are calculated by numerically solving the flux balance equation,

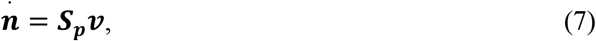

under the constraint that number of each intermediate chemical species does not change over a reaction cycle, and that number of target molecules increases by 1,

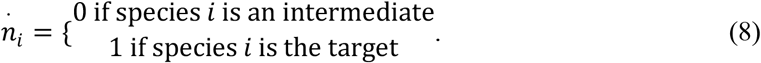

The calculated stoichiometries for synthesis of each branched chain hydrocarbon considered in this article are listed in **Table S2**. The stoichiometries for each molecule are then combined with their molecular weights and energies per molecule (listed in **Table S3**) to calculate the energy input and production efficiency (FIG-4A&B.PY and FIG-4A&B.PY, results shown in **Fig. 4**). The energy inputs and conversion efficiencies for jet fuel blends are calculated by a weighted average of the energy inputs and efficiencies of the individual components (**Fig. 5**).

**Figure 5.**
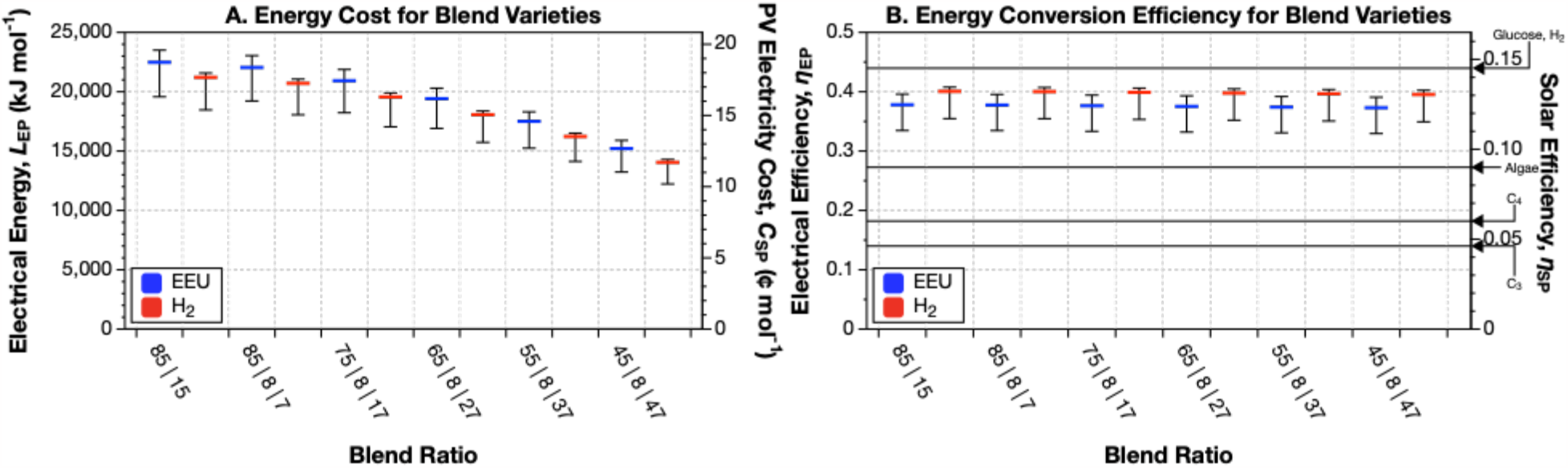
Effect of adding a proportion of branched-chain hydrocarbons to existing fuel blend estimates lowers efficiencies by only ∼.1% for every 10% branched chains. Effect of changing carbon-fixation method on **(A)** energy input for, and **(B)** energy conversion efficiency of production of a model jet fuel blend containing 85% molarity (equal numbers of C_10_ to C_16_ molecules) and 15% terpenoids (equal numbers of each of the 5 terpenoids) when using the ADO alkane termination pathway. The central value (thick blue or red bar) corresponds to the most likely value of the trans-membrane (Δ*U*_membrane_) voltage of 140 mV. Meanwhile, Δ*U*_membrane_ = 280 mV produces lower efficiencies (hence a higher energy input), while Δ*U*_membrane_ = 80 mV produces higher efficiencies (and hence lower energy inputs)^8^. This figure can be reproduced by running the codes INFO-FIG-5A&B.PY, and FIG-5A&B.PY in the EMP-TO-BRANCHED-JET online code repository^39^.

### Restrictions of Branched Chain Formation

Due to the limits of FAS, we are unable to produce adjacently-methylated branched-chain alkanes. Given the molecular structure of malonyl-CoA, a methyl group can only be added to every second carbon during lengthening. This prevents the preparation of some single-methyl and multi-methylated branched-chains. Nonetheless, FAS allows us to produce a great variety of compounds that allow us to more closely mimic the composition of traditional jet fuel blends.

### Production of (S)-methyl-malonyl-[acp] Lengthener

Branches are added to a growing alkane during lengthening by incorporation of a (S)-methyl-malonyl-[acp]. (S)-methyl-malonyl-CoA is produced by a side reaction of Acetyl-CoA Carboxylase (ACACA). Typically ACACA reacts acetyl-CoA with HCO_3_^-^ to produce malonyl-CoA. However, occasionally propionyl-CoA is used in place of acetyl-CoA place, generating (S)-methyl-malonyl-CoA. Under normal circumstances the cell degrades (S)-methyl-malonyl-CoA back to propionyl-CoA by methylmalonyl-CoA decarboxylase (MMCD)^3^ [Linster2011a]. But, down-regulation of MMCD leads to the accumulation of (S)-methyl-malonyl-[acp], and thus the production of branched chain fatty acids^3^.

## Results and Discussion

Herein, we calculate conversion efficiencies of a panel of 13 single-branched and 11 multi-branched hydrocarbons with backbone lengths between C_5_ and C_10_, using H_2_-oxidation or EEU for electron delivery. **Figs. 4A** and **4C** show the energy required to produce a mole of each hydrocarbon, and **Figs. 4B** and **4D** show the electrical-and solar-to-chemical conversion efficiency for each molecule. **Fig. 5** shows the energy costs and production efficiencies of jet fuel blends containing increasing amounts of branched chain alkanes.

### Electron Uptake by H_2_-Oxidation Produces 2% Higher Energy Conversion Efficiencies than EEU

As seen in previous studies, H_2_-oxidation is a more efficient method for electron delivery than EEU^4,8,25,26^. For single-and multi-methylated branched chain hydrocarbons, use of H_2_-oxidation raises the electricity-to-fuel energy conversion efficiency by ≈ 2.3%. The energy-cost-savings of H_2_-mediated EMP over EEU-mediated EMP of single-branched-chain hydrocarbons ranges from 567 kJ mol^-1^ for 3-methyl-pentane (3-M_1_-pentane) to 1,008 kJ mol^-1^ for 5-methyl-decane (5-M_1_-decane). For multi-methylated branched-chain hydrocarbons, the energy savings range from 886 kJ mol^-1^ for 3,5-dimethyl-octane (3,5-M_2_-octane) to 1,252 kJ mol^-1^ for 2,4,6,8-tetramethyl-decane (2,4,6,8-M_4_-decane).

### Electromicrobial Production Could Achieve Synthesis of Single-branched-chain Alkanes at Efficiencies Between 34.7% and 40.0%

The energy requirements for synthesis of single-branched-chain alkanes range from 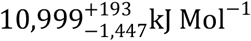 for H_2_-driven 3-M_1_-pentane production to 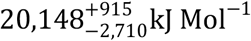 for EEU-driven 5-M_1_-decane (**Fig. 4A**). We observe a general trend of increased energy cost with increased chain length. However, changing the position of the methylation sites can change the energy cost of synthesis. For example, the production of 3-M_1_-hexane is more expensive than 2-M_1_-hexane (see **Fig. 2C**). This difference in energy is due to the higher energy cost for production of the (S)-methyl-malonyl-[acp] lengthener needed to install the branch in 3-M_1_-hexane, versus the cost of the 2-methyl-propionyl-[acp] initiator needed to install the branch in 2-M_1_-hexane. The high cost of (S)-methyl-malonyl-[acp], which adds 2 carbons to backbone, is due to the high energy cost of synthesis (15 ATP and 7 NADH, or 7.5 ATP C^-1^ and 3.5 NADH C^-1^). In contrast, 2-methyl-propionyl-[acp] adds 3 carbons to the backbone and requires 14 ATP and 10 NADH (4.6 ATP C^-1^ and 3.3 NADH C^-1^).

The electrical-to-fuel energy conversion efficiencies of single-branched-chain alkane production range from 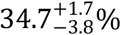 for 2-M_1_-pentane with EEU, to 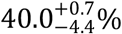 for 2-M_1_-octane with H_2_ (**Fig. 4B**). All of these efficiencies are close to the production efficiency for butanol (44% for the H_2_-driven Calvin cycle^8^) and glucose (44.6% for the H_2_-driven Calvin cycle^25^).

The production efficiency of odd-length alkanes with a branch on the second carbon is lower than that for even-length alkanes with the branch in the same place. Odd-alkanes with a branch on the second carbon need to be initiated with energy-expensive 3-methyl-butanoyl-[acp]. On the other hand, even-length alkanes with the branch in the same place need to be initiated with energy-cheap 2-methyl-propionyl-[acp] (see 2-M_1_-pentane and 2-M_1_-hexane an example in **Fig. 2C**). Synthesis of 2-methyl-propionyl-[acp] in total costs 14 ATP and 10 NADH and adds 3 carbons to backbone (4.6 ATP C^-1^ and 3.3 NADH C^-1^). In contrast, 3-methyl-butanoyl-[acp] adds 4 carbons to the backbone, but requires 21 ATP and 14 NADH (5.25 ATP C^-1^ and 3.5 NADH C^-1^).

The efficiency of production of straight-chain alkanes ranges from 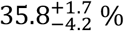 for hexane to 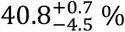 for hexadecane^4^. Though similar to the efficiencies of our branched-chains, our straight-chains appear to have slightly higher efficiencies across the board, likely a result of higher combustion energies for compounds of similar carbon length. In all cases, if the electricity for production of these alkanes is derived from a perfectly efficient solar photovoltaic^40^, then their production efficiency exceeds the efficiency of all forms of photosynthesis (see the right hand axis in **Fig. 4B**).

### Electromicrobial Production Could Achieve Synthesis of Multi-branch-chain Alkanes with Efficiencies Between 35.6% and 39.9%

The energy costs of electromicrobial production of multi-branched chain alkanes range from 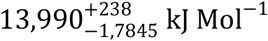 Mol^−1^ for H_2_-derived 2,4-M_2_-hexane to 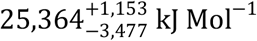 Mol^−1^ for 2,4,6,8-M_4_-decane produced with EEU (**Fig. 4C**). As with single-branched chain alkanes there is a general increase in energy requirement with hydrocarbon length. Furthermore, as with single-branched chain alkanes, the choice of initiator molecule and the sites of branching cause notable differences in energy cost. The drop of energy required for 3,5-dimethyl octane over 2,4-dimethyl octane can be attributed to this cause. Here, 3,5-dimethyl octane obtains its branches entirely from the use of (S)-methylmalonyl-[acp], which is energetically unfavorable given its use of propionyl-[acp] in production. Contrastingly, 2,4-M_2_-octane requires the initial production of 2-methyl-propionyl-[acp], and further utilization of (S)-methylmalonyl-[acp]. 2-methyl-propionyl-[acp] is less energetically expensive to produce and thus costs less than (S)-methylmalonyl-[acp] alone. Therefore, though both molecules are chemically similar and combust similarly, they differ in production cost by ∼600 kJ Mol^−1^.

The electrical-to-fuel energy conversion efficiencies for multi-branched chain alkanes are similar to those for single-branched-chain alkanes, ranging from 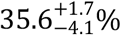 for 2,5-M_2_-heptane with EEU to 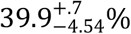 for 2,4-M_2_-decane with H_2_ (**Fig. 4D**).

### Increasing the Fraction of Branched-chain Hydrocarbons in a Jet Fuel Blend by 10% Lowers Conversion Efficiency by 0.1%

We next calculated how the introduction of branched-chain alkanes into a jet fuel blend would change the energy production costs and energy conversion efficiency (**Fig. 5**). We compared a previously conceived blend containing 85% straight-chain alkanes (C_10_-C_16_) and 15% terpenoids (pinene, limonene, farnesene, bisabolene, and geraniol)^4^, with two additional blends incorporating our branched chain hydrocarbons (C_8_-C_10_ backbone). The energy-conversion efficiency for H_2_-driven production of original blend is 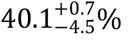.

The first new blend contains 7% branched-chain alkanes, a minimum-allowed 8% terpenoids^41^, and is filled out with 85% C_10_ to C_16_ straight-chain alkanes. The conversion efficiency for H_2_-driven production of this blend is slightly decreased to 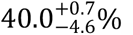. The energy conversion efficiency for H_2_-driven production a 17% branched-chain alkanes, 8% terpenoid, and 75% straight-chain alkanes blend is further decreased to 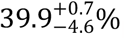. As we further increase branched-chain content, we see a continuation of this trend, with a drop in energy conversion efficiency of 0.1% for every additional 10% of branched-chain content.

## Conclusion

In this article we calculated the energy costs and energy-to-fuel conversion efficiencies of electromicrobial production of individual branched-chain alkanes and fuel blends containing them. When using H_2_-oxidation for electron delivery, the Calvin cycle for CO_2_-fixation, and the ADO pathway for decarboxylation we were able to see efficiencies as high as 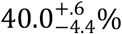 for single-branched-chain 3-M_1_-hexane and 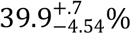 for multi-branched-chain 2,4-M_2_-decane (**Fig. 4**). When supplied with solar electricity, these efficiencies far exceed those of all forms of photosynthesis.

Replacing terpenoids with branched-chain alkanes in a jet fuel blend increases the electromicrobial production efficiency. A previously developed 85% straight-chain alkane and 15% terpenoid blend can be produced with 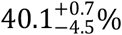 electrical-to-fuel efficiency, which decreases steadily with branched-chain content at a rate of .1% per 10% total branched-chain content.

Making this work a reality will require extensive metabolic engineering and synthetic biology, concerning both the creation of novel and potentially toxic pathways for producing hydrocarbons in combination with the engineering of radical new hosts for EMP. However, we have established that the high theoretical efficiency of EMP justifies doing this arduous engineering work in the hopes of creating viable, sustainable biofuels for demanding applications like aviation. Further, by using EMP to create a library of diverse, branched hydrocarbons which go beyond simple unbranched alkanes, we can create a repository of fuel components which when blended can replicate the desirable attribute of today’s fuels, furthering the cause of biofuels ultimately sourced from renewable electricity.

## End Notes

### Code Availability

All code used in calculations in this article is available at https://github.com/barstowlab/emp-to-branched-jet and is archived on Zenodo^39^.

### Materials & Correspondence

Correspondence and material requests should be addressed to B.B..

## Author Contributions

Conceptualization, T.J.S., D.A.S. and B.B.; Methodology, T.J.S., D.A.S. and B.B.; Investigation, T.J.S. and D.A.S.; Writing - Original Draft, T.J.S. and D.A.S.; Writing - Review and Editing, T.J.S., D.A.S. and B.B.; Resources, B.B.; Supervision, B.B..

## Acknowledgements

This work was supported by a Cornell Energy Systems Institute Postdoctoral Fellowship to D.A.S.; Cornell University startup funds, a Career Award at the Scientific Interface from the Burroughs-Wellcome Fund, U.S. Department of Energy Biological and Environmental Research grant DE-SC0020179, a Cornell 2030 Project Fast Grant, and a gift from Mary Fernando-Conrad and Tony Conrad to B.B..

## Competing Interests

The authors declare no competing interests.

## Notes

https://doi.org/10.5281/zenodo.7693794

### Competing Interest Statement

The authors have declared no competing interest.

