## Supplementary Information for "Efficiency Estimates for Electromicrobial Production of Branched-chain Hydrocarbons"

†Corresponding author:

### Supplementary Information Tables

**Table S1.** Symbols used in this article.

**Table S2.** NAD(P)H, reduced Ferredoxin, and ATP needed for synthesis of branched-chain alkanes.

**Table S3.** Molecular weights and energy densities for branched-chain alkanes.

| Symbol | Unit | Description |
|---|---|---|
| $E_{HC}$ | J molecule$^{-1}$ | Energy carried per hydrocarbon molecule. |
| $\dot{N}_{HC}$ | molecule s$^{-1}$ | Hydrocarbon molecules produced per second. |
| $N_A$ | molecule mol$^{-1}$ | Avogadro constant. |
| $P_{e,\,T}$ | J s$^{-1}$ | Total electrical power input into electromicrobial production system. |
| $L_{EP}$ | kJ mol$^{-1}$ | Electrical energy cost to generate one mole of product. |
| $C_{SP}$ | ¢ mol$^{-1}$ | Minimum solar electricity cost for synthesis of one mole of product at 2030 solar electricity [illegible] |
| $M_{HC}$ | g mol$^{-1}$ | Molecular weight of hydrocarbon molecule. |
| $\eta_{EP}$ | % | Electrical to product (*e.g.,* hydrocarbon) energy conversion efficiency. |
| $\eta_{SP}$ | % | Solar to product (*e.g.,* hydrocarbon) energy conversion efficiency. |
| $e$ | A s | Fundamental charge. |
| $\nu_{ep}$ | # | Number of electrons needed for synthesis of a product (*e.g.*, hydrocarbon) molecule. |
| $\Delta U_{e.cell}$ | V | Potential difference across bio-electrochemical cell. |
| $\xi_{I2}$ | # | Faradaic efficiency of the bio-electrochemical cell. |
| $\nu_{p,\,NADH}$ | # | Number of NAD(P)H molecules needed to make a final product molecule. |
| $\nu_{p,\,Fd}$ | # | Number of Fd molecules needed to make a final product molecule. |
| $\nu_{p,\,ATP}$ | # | Number of ATP molecules needed to make a final product molecule. |
| $\Delta G_{ATP/ADP}$ | J | Free energy for regeneration of ATP. |
| $\Delta U_{membrane}$ | V | Inner membrane potential difference. |
| $U_{H2}$ | V | Standard potential of proton reduction to $H_2$. |
| $U_{acceptor}$ | V | Standard potential of terminal electron acceptor reduction. |
| $U_Q$ | V | Redox potential of the inner membrane electron carrier. |
| $U_{NADH}$ | V | Standard potential of NADH. |
| $U_{Fd}$ | V | Standard potential of Ferredoxin. |

**Table S1.** Symbols used in this article.

| **Branched-chain Alkane** | **No. ATP** | **No. NAD(P)H** | **No. Reduced Ferredoxin** | **No. $CO_2$** |
|---|---|---|---|---|
| 2-$M_1$-pentane | 29 | 22 | 0 | 6 |
| 2-$M_1$-hexane | 31 | 24 | 0 | 7 |
| 2-$M_1$-heptane | 37 | 28 | 0 | 8 |
| 2-$M_1$-octane | 39 | 30 | 0 | 9 |
| 3-$M_1$-pentane | 30 | 21 | 0 | 6 |
| 3-$M_1$-hexane | 37 | 24 | 0 | 7 |
| 3-$M_1$-heptane | 38 | 27 | 0 | 8 |
| 3-$M_1$-octane | 45 | 30 | 0 | 9 |
| 3-$M_1$-nonane | 46 | 33 | 0 | 10 |
| 4-$M_1$-octane | 45 | 30 | 0 | 8 |
| 4-$M_1$-decane | 53 | 36 | 0 | 11 |
| 5-$M_1$-nonane | 46 | 33 | 0 | 10 |
| 5-$M_1$-decane | 53 | 36 | 0 | 11 |
| 2,4-$M_2$-hexane | 37 | 27 | 0 | 8 |
| 2,4-$M_2$-octane | 46 | 33 | 0 | 10 |
| 2,4-$M_2$-decane | 54 | 39 | 0 | 12 |
| 2,5-$M_2$-heptane | 44 | 31 | 0 | 9 |
| 2,5-$M_2$-nonane | 52 | 37 | 0 | 11 |
| 3,5-$M_2$-octane | 552 | 33 | 0 | 10 |
| 3,5-$M_2$-decane | 60 | 39 | 0 | 12 |
| 4,6-$M_2$-decane | 60 | 39 | 0 | 12 |
| 2,4,6-$M_3$-octane | 53 | 36 | 0 | 11 |
| 2,4,6-$M_3$-decane | 61 | 42 | 0 | 13 |
| 2,4,6,8-$M_4$-decane | 68 | 45 | 0 | 14 |

**Table S2.** NAD(P)H, reduced Ferredoxin, and ATP needed for synthesis of single molecules of branched-chain alkanes from $CO_2$. All alkanes are synthesized from $CO_2$ with the Calvin cycle, Type II Fatty Acid Synthesis and with the ADO decarboxylation pathway. Calculated with the INFO-FIG4A&B.PY, INFO-FIG4C&D.PY codes in the EMP-TO-BRANCHED-JET repository [Sheppard2023b].

| Branched-chain Alkane | Molecular Weight (Da) | Energy Density (kJ mol$^{-1}$) | Energy Density (J molecule$^{-1}$) |
|---|---|---|---|
| 2-$M_1$-pentane | 86.18 | 4,153.80 | 6.90E-18 |
| 2-$M_1$-hexane | 100.2 | 4,809.98 | 7.99E-18 |
| 2-$M_1$-heptane | 114.23 | 5,466.16 | 9.08E-18 |
| 2-$M_1$-octane | 128.26 | 6,140.40 | 1.02E-17 |
| 3-$M_1$-pentane | 86.18 | 4,159.82 | 6.91E-18 |
| 3-$M_1$-hexane | 100.2 | 4,816.00 | 8.00E-18 |
| 3-$M_1$-heptane | 114.23 | 5,478.20 | 9.10E-18 |
| 3-$M_1$-octane | 128.26 | 6,140.40 | 1.02E-17 |
| 3-$M_1$-nonane | 142.28 | 6,742.40 | 1.12E-17 |
| 4-$M_1$-octane | 128.26 | 6,140.40 | 1.02E-17 |
| 4-$M_1$-decane | 156.31 | 7,404.60 | 1.23E-17 |
| 5-$M_1$-nonane | 142.28 | 6,742.40 | 1.12E-17 |
| 5-$M_1$-decane | 156.31 | 7,404.60 | 1.23E-17 |
| 2,4-$M_2$-hexane | 114.23 | 5,461.69 | 9.07E-18 |
| 2,4-$M_2$-octane | 142.28 | 6,768.13 | 1.12E-17 |
| 2,4-$M_2$-decane | 170.33 | 8,074.58 | 1.34E-17 |
| 2,5-$M_2$-heptane | 128.26 | 6,114.91 | 1.02E-17 |
| 2,5-$M_2$-nonane | 156.31 | 7,421.35 | 1.23E-17 |
| 3,5-$M_2$-octane | 142.28 | 6,768.13 | 1.12E-17 |
| 3,5-$M_2$-decane | 170.33 | 8,074.58 | 1.34E-17 |
| 4,6-$M_2$-decane | 170.33 | 8,074.58 | 1.34E-17 |
| 2,4,6-$M_3$-octane | 156.31 | 7,421.35 | 1.23E-17 |
| 2,4,6-$M_3$-decane | 184.36 | 8,727.80 | 1.45E-17 |
| 2,4,6,8-$M_4$-decane | 198.39 | 9,381.02 | 1.56E-17 |

**Table S3.** Molecular weights and energy densities for branched-chain alkanes. Data from NIST database [NIST2022a].

### Supplementary Information References

[NIST2022a] P.J. Linstrom and W.G. Mallard, Eds., NIST Chemistry WebBook, NIST Standard Reference Database Number 69, National Institute of Standards and Technology, Gaithersburg MD, 20899 (retrieved September 24, 2022). doi:10.18434/T4D303.

[Sheppard2023b] T.J. Sheppard and B. Barstow,
